# Psychophysiological stress induced neurobehavioral changes in adolescent rat and their impact in Adult rat enhanced by cinnamaldehyde

**DOI:** 10.1101/2022.08.14.503911

**Authors:** C S Keerthi Priya, R Ravindran

## Abstract

Adolescent is a tricky developmental period were the Physical, cognitive and social changes takes place. Adolescent brain undergoes vigorous maturation and more susceptible to stressors, distresses in the brain development leading to more susceptibility for stress related psychophysiological disorders in later life. The present study explores the association of adolescent stress on same period and on adult. Adolescent rats were restraint to induce psychophysiological stress from PND28 to PND42.Cinnamaldehyde (CA) 25 mg/kg body weight is treated along with the stress. Restraint stress showed more deleterious effect on learning, memory, anxiety, depression and locomotor activity. The result of this study reveal that adolescent stress has impact on that period and even long-lasting impact. But CA may act as antistressor and cognitive enhancer.

## INTRODUCTION

Early life stress, solely experienced at the first retro of life, distresses the brain developmental trajectories leading to improved susceptibility for stress related psychophysiological disorders later in life. Now a day’s stress is not limited only to adults, but also to adolescence and have a profound long-lasting effect on a number of behavioral and neural outcomes impact on the health and psychological functioning of the individual. Brain structure and function shows a remarkable degree of plasticity during adolescence[1]. It is believed that adolescence is a period of increased vulnerability and that stress-induced perturbations in the developing adolescent brain may contribute to altered functioning later in life. Exposure to stress during adolescence alters stress reactivity[2], cognitive ability, emotionality[3], and hippocampal structure in adulthood.

The effects of stress during adolescence may be permanent, unlike in adulthood where many effects of stress on the hippocampus are reversible with time[3].Cinnamaldehyde **(CA)**(3-phenyl-2-propenal) has been in public use since 1900. CA is a light yellowish fluid with a sweet, spicy, warm odor, and a pungent taste, which is indicative of cinnamon[4].CA have many pharmacological activities such as anti-diabetic, antioxidant, cognitive enhancer, anti-inflammatory, anti-cancer activity, suppresses neuronal apoptosis, glial activation, and Aβ burden in the hippocampus and protects memory and learning [5].

## AIM

This study is aim to investigate whether a stressful experience during adolescence postnatal day (**PND 28 - 42**) effects are observed in both adolescence (PND42) adult (**PND65**) neurobehavioral alterations and effect of cinnamaldehyde (**CA**) in stress retrieval.

## MATERIALS AND METHOD

### Animals

Postnatal 28days (PND28) Wistar albino male rats weighing 70-80 g was used for the study. Animals were housed in a group and maintained in controlled room temperature 23°C±2°C with 12:12-h light: dark cycle and allowed to free access to food and water. All animal procedures were approved by the Institutional Animal Ethical Committee and CPCSEA (IAEC No: 03/02/19). All efforts were made to minimize both the number of animals used and unwanted suffering to the animals during experimental procedures.

### Experimental design

Animals were divided into four group; each group consists of six animals: Group I - Control, Group II –Restraint stress, Group III-Treatment alone (CA), Group IV-Restraint stress+ Treatment (CA). To avoid circadian rhythm-induced variation, all the experiments have been carried out between 9 and 10 AM.

### Psychophysiological Stress Model (PS)

The animal model of psychophysiological stress was induced by restrainer for 6hrs/15 days[6]. It is made of cylindrical plexiglass tubes[7] with small holes at the bottom for aeration and for excretion of urine and fecal matter. Many tubes with different size range to accommodate growing rat till PND42.They were also food and water deprived during the stress induction period. Cylindrical plexiglass tubes with animals during the stress exposure are kept adjacent to circumvent isolation stress.

### Drug

Cinnamaldehyde (CA) 25mg/kg body weight[8] is mixed with 1% TWEEN 80 and sterile saline are used as a vehicle[9] administrated orally for 15 days before start of the psychophysiological stress. CA was purchased from Sigma-Aldrich (**W228613**) is used for the study.

### Overt Behavior recordings and analysis

Behavior was recorded using a portable camera fed to a laptop and analyzed using a tracking and video analyzer software, ANY-maze version 6.18(New Delhi, INDIA).

All the results were analyzed by **TWO WAY ANOVA** using **Graph pad prism**. Post test by Bonferroni. **P** value > 0.01 is considered as significant. Results were designated significant when the P-value (P) < 0.05: *= P < 0.05, **= P < 0.01, ***= P < 0.001, **** = P < 0.0001, ns = non-significant.

### Neurobehavioral studies

#### 1. Learning and Memory

##### 1.1. Eight Arm Radial Maze

Spatial learning and memory was tested by using an eight arm radial maze[10].The eight arm radial maze made of steel material, had an octagonal central platform, 33.5 cm wide, around which were arranged 60 cm long by 12 cm wide arms. The whole apparatus was elevated 40 cm from the floor in a sound proof chamber.

Each individual rat had its own set of four rewarded arms (sucrose pellets). The room contained several visual reference cues on the wall. Each trial began with the placement of the animal on the central platform facing arm number one and ended when the rat had visited the four baited arms or after a period of 5 min. The following parameters were measured[11].

###### Based on Olton’s definition

1. Number of reference memory errors, (i.e. each entry into a non-baited arm)
2. Number of working memory errors, (i.e. re-entries into already visited baited arms were noted)
3. Time taken to visit all the baited arms.

##### 1.2. Novel Object Recognition Task

The task is used to analyze recognition memory[12]. The 1^st^ day animals were habituated to the open-field apparatus. 2nd day (Trail 1 or T1), animals were familiarized with identical objects placed diagonally opposite in the central squares of the open field (ball of same size and shape) for 5 min. After 2 h of time interval (Trail 2 or T2), one familiar object is replaced with a novel object (Multi color cube) and now the time spent in exploring the objects was recorded. Percentage of time spent was calculated as **TNovel/(TNovel+TFamiliar) ×100,** where T Novel is time spent with a novel object and TFamiliar is time spent with a familiar object.

#### 2. Anxiety and Depression

##### 2.1. Place preference task

Place preference task (PPT) is an approach avoidance conflict between the novel environment and evasion of brightly lit, open space[13]. The box was divided into two compartments, 18 × 15 × 15 inches (long, wide and high) light compartment with open at the top and 12 × 15 × 15 inches (long, wide and high) the dark compartment that was fully enclosed. The divider between the two compartments and contained a 3 × 4 inch (wide, high) opening at floor level. This allows the animal entries between compartments. The measures scored were (1) initial latency to enter the dark compartment, (2) time spent in brighter area and (3) time spent in the dark compartment[14].

##### 2.2 Hole Board task

It consists of wooden, black box with white lines in the surface measuring 68 cm x 68 cm. it is 40 cm high, and box was raised 28 cm with four holes (4cm in diameter). Apparatus was placed above the ground on a metal stand. Room is well illuminated and clues are placed. At the end of the each trail the apparatus was cleaned with 70% alcohol. It is mainly to access anxiety and exploratory like behavior,

##### 2.3 Elevated Plus Maze

The elevated plus maze (EPM) is used to assess innate anxiety in rodents. The EPM apparatus made of wood consists of a “+”-shaped maze elevated above the floor with two oppositely positioned closed arms, two oppositely positioned open arms, and a center area[15]. No pre exposure to apparatus, they were tested directly Rats were placed in the central square of the maze, facing one of the open arms. The tested area was cleaned with 70% alcohol prior to the introduction of each animal. Animals falling off the maze were eliminated from the analysis. The parameters include 1, number of open arm entries,2. closed arm entries and the 3. number of head dips (dipping the head below the open arm)[16].

#### 3. Socialbility and locomotor activity

##### 3.1. Three Chamber Test (TCT)

The Three-Chamber test (TCT) assesses cognition in the form of general sociability and interest in social novelty in rodent models. The apparatus comprised of three chambers rectangular box, a center compartment of (20 cm × 35 cm × 35 cm) with a left and a right compartment of (30 cm × 35 cm × 35 cm) fabricated in plexiglass material. The dividing walls had retractable doorways allowing access to each chamber and two jail like cup cages were 10 cm in height, with a bottom of 9 cm and bars placed 1 cm gap with tightly closed cover to hold the stranger animals on each side of the chamber. The parameters accessed are time spent with the familiar animal and time spent with the unfamiliar animal. This test is useful for quantifying deficits in social behavior in rodents[17]

##### 3.2. Open Field test

Open field task is used to measure the exploratory and anxiety related behavior[18] in both adolescent (PND43) and adult (PND 65) rats. The apparatus size was altered according to the age of rat (80cmx80cm)[19], (100×100 cm) for adolescents and adult rats respectively. The apparatus was dived into 25 equal squares and was cleaned after the end of each animal testing with 70% alcohol to remove olfactory clues. The task was carried out without any exposure to the apparatus one day after the end of the stress with clues in the four side of the wall in the room and was assessed by the following parameters such as 1.Peripheral ambulation(no of entry),2. Central ambulation(no of entry),3. Rearing (standing on the hind limbs and sometimes leaning on the wall with forelegs, sniffing and looking around),4.Grooming (licking the fur, washing face or scratching behavior), 5. Immobilization (no activity of the animal) and 6.Defection (number of fecal pellets)for 5 mins.

#### 4. Cognitive Behavior

##### 4.1. Attention set shifting task (ASST)

The attentional set shifting task (ASST) is used to assess cognitive flexibility including reversal learning and memory. Apparatus (70 x 40 x 18 cm) consists of two chambers separated by a plexiglass in which a small door is present for the entry of the rat from the waiting area to the testing area. Testing chamber is equally divided in between the two pots (7 x 4 cm) to avoid hinderance to the rat. Animals were partially fasted. In the habituation phase one of the bowl was baited, so that rat could move to the unbaited bowl and learn the cue contingency. Trials are continued until six successive attempts are made. The following are the parameters **Simple Discrimination** (**SD**) only relevant dimension present (i.e., it could be odor). **Compound Discrimination (CD) introduces** second dimension (i.e., medium). The **first reversal (Rev 1)** would require the mouse to respond in the bowl with the previously irrelevant odor. Intradimenstional **(ID)** are new compound stimuli, differing again according to odor and digging medium, are introduced. Reverse of ID is (**Rev 2**). Extradimensional (**ED**) the exemplars are changed again, but now the relevant dimension in incongruent with the prior stages. Reverse of ED is (**Rev 2**).Bowl is baited with different odor and medium. Odor is blotted in the blotting paper which is digged in the medium without giving clue to the rats.

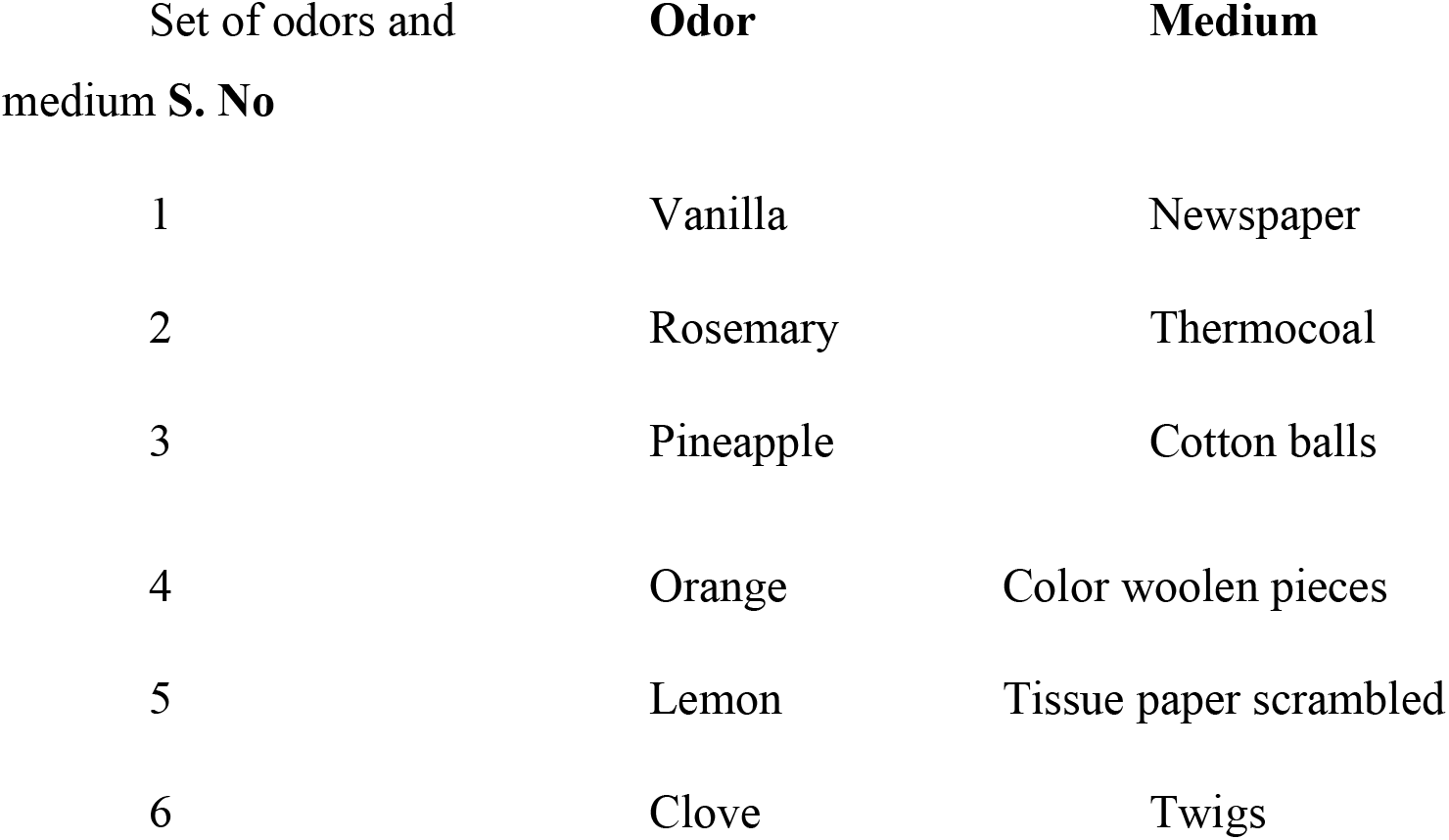

| Set of odors and medium S. No | Odor | Medium |
| --- | --- | --- |
| 1 | Vanilla | Newspaper |
| 2 | Rosemary | Thermocoal |
| 3 | Pineapple | Cotton balls |
| 4 | Orange | Color woolen pieces |
| 5 | Lemon | Tissue paper scrambled |
| 6 | Clove | Twigs |

## RESULTS

### Learning and memory

Working Memory Error were markedly varying among the groups. The increased WME were observed in RS PND42 & PND65 (2.167±0.477 & 1.667±0.333) when compared to control (0.333±0.211 &0.333±0.211) P value is < 0.001 & P<0.01. WME decreased in RS+T (0.833±0.307,0.833±0.307) and p value is <10.001 & p<0.01 group when compared to RS.

Reference Memory Error were markedly varying among the groups. The increased RME were observed in RS PND42 & PND65 (3.667±0.667&2.500±0.428) when compared to control (0.667±0.211 & 0.500±0.224) P value is < 0.001 & P<0.01. RME decreased in RS+T (1.333±0.422 & 1.333±0.422) and p value is p<0.001 & p>0.05 group when compared to RS.

Time Taken is increased in RS PND42 & PND65 (59.66±3.844 & 63.500±3.423) when compared to control (28.667±1.022 & 31.667±1.892) P value is < 0.01 & P < 0.001. Time Taken is decreased in RS+T (36.667±1.520 & 31.883±1.641) and p value is p<0.001 & p<0.001 group when compared to RS.

Novel object location was markedly varying among the groups. The decreased object location was observed in RS PND42 & PND65 (49.500±1.670 & 41.000±2.463) when compared to control (70.167±1.352 & 69.000±0.856) P value is < 0.001 & P<0.001. Object location increased in RS+T (60.167±1.352 & 55.833±1.537) and p value is p<0.001 & p<0.001 group when compared to RS **Table 2 A**.

**Table 1.** Mean± SEM for RAM. where WME is working memory error, RME is Reference memory error and TT is time taken.

| GROUPS | CON | RS | RS+T | T |
| --- | --- | --- | --- | --- |
| PND42(WME) | $0.333 \pm 0.211$ | $2.167 \pm 0.477$ | $0.833 \pm 0.307$ | $0.833 \pm 0.307$ |
| PND65(WME) | $0.333 \pm 0.211$ | $1.667 \pm 0.333$ | $0.833 \pm 0.307$ | $0.500 \pm 0.224$ |
| PND42(RME) | $0.667 \pm 0.211$ | $3.667 \pm 0.667$ | $1.333 \pm 0.422$ | $0.833 \pm 0.307$ |
| PND65(RME) | $0.500 \pm 0.224$ | $2.500 \pm 0.428$ | $1.333 \pm 0.422$ | $0.833 \pm 0.307$ |
| PND42(TT) | $28.66 \pm 1.022$ | $59.66 \pm 3.844$ | $36.667 \pm 1.520$ | $28.833 \pm 0.401$ |
| PND65(TT) | $31.667 \pm 1.892$ | $63.500 \pm 3.423$ | $31.883 \pm 1.641$ | $30.000 \pm 1.155$ |

**Table 2 A.**
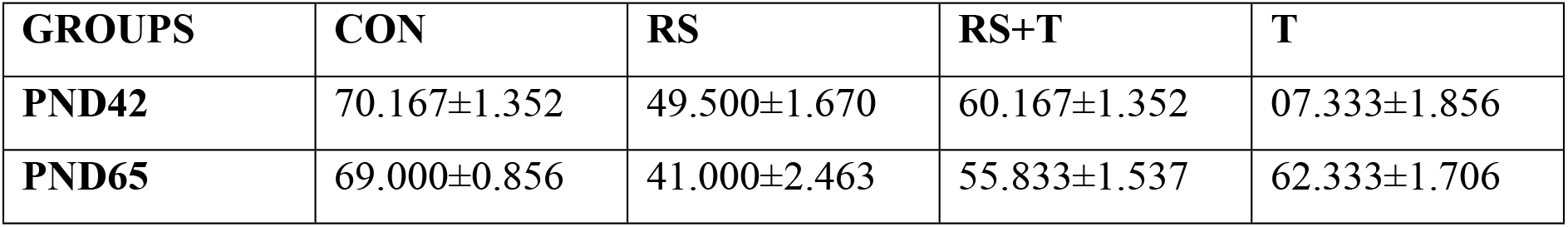
Mean ± SEM for Novel object location task

| GROUPS | CON | RS | RS+T | T |
| --- | --- | --- | --- | --- |
| PND42 | 70.167±1.352 | 49.500±1.670 | 60.167±1.352 | 07.333±1.856 |
| PND65 | 69.000±0.856 | 41.000±2.463 | 55.833±1.537 | 62.333±1.706 |

**Table 2 B.** Mean ± SEM for Novel object recognition task

| GROUPS | CON | RS | RS+T | T |
| --- | --- | --- | --- | --- |
| PND42 | 64.833±1.740 | 29.500±1.335 | 42.500±2.141 | 60.167±1.424 |
| PND65 | 72.000±1.095 | 33.167±1.537 | 41.833±2.469 | 55.833±1.537 |

Novel object recognition was markedly varying among the groups. The decreased object recognition was observed in RS PND42 & PND65 (29.500±1.335 & 33.167±1.537) when compared to control (64.833±1.740 & 72.000±1.095) P value is < 0.001 & P<0.001. Object recognition increased in RS+T (42.500±2.141 & 41.833±2.469) and p value is p<0.01 & p<0.01 group when compared to RS **Table 2 B**.

#### 2. Anxiety and Depression

Time taken in brighter area was markedly varying among the groups. The decreased time spent was observed in RS PND42 & PND65 (93.770±6.953 & 72.383±8.882) when compared to control (188.893±4.472 & 189.333±4.951) P value is < 0.001 & P<0.001. TT BA increased in RS+T (137.167±10.117 & 93.027±1.792) and p value is p<0.001 & p<0.001 group when compared to RS **Table 3**.

**Table 3.** Mean± SEM for Place preference Task where TT BA is Time taken in Bright area, TT DA is Time taken in dark area, N FB is number of fecal bolus.

| GROUPS | CON | RS | RS+T | T |
| --- | --- | --- | --- | --- |
| PND42(TT BA) | $188.893 \pm 4.472$ | $93.770 \pm 6.953$ | $137.167 \pm 10.117$ | $183.303 \pm 1.565$ |
| PND65(TT BA) | $189.333 \pm 4.951$ | $72.383 \pm 8.882$ | $93.027 \pm 1.792$ | $182.628 \pm 1.356$ |
| PND42(TT DA) | $110.962 \pm 4.404$ | $210.680 \pm 3.549$ | $162.833 \pm 10.117$ | $116.697 \pm 1.565$ |
| PND65(TT DA) | $110.667 \pm 4.951$ | $227.617 \pm 8.882$ | $206.973 \pm 1.792$ | $117.372 \pm 1.356$ |
| PND42(N FB) | $2.167 \pm 0.401$ | $3.333 \pm 0.211$ | $2.500 \pm 0.224$ | $2.000 \pm 0.365$ |
| PND65(N FB) | $1.833 \pm 0.307$ | $2.500 \pm 0.224$ | $1.500 \pm 0.224$ | $0.667 \pm 0.211$ |

Time taken in dark area was markedly varying among the groups. The decreased time spent was observed in RS PND42 & PND65 (93.770±6.953 & 72.383±8.882) when compared to control (188.893±4.472 & 189.333±4.951) P value is < 0.001 & P<0.001. TT BA increased decreased in RS+T (137.167±10.117 & 93.027±1.792) and p value is p<0.001 & p<0.05 group when compared to RS **Table 3**.

Fecal bolus was markedly varying among the groups. The increased number of fecal bolus was observed in RS PND42 & PND65 (3.333±0.211 & 2.500±0.224) when compared to control (2.167±0.401 & 1.833±0.307) P value is < 0.05 & P<0.05. Number of fecal bolus decreased in RS+T (2.500±0.224 & 1.500±0.224) and p value is p>0.05 & p<0.05 group when compared to RS **Table 3**.

Dipping was markedly varying among the groups. The decreased number of dipping was observed in RS PND42 & PND65 (3.333±0.211 & 0.333±0.211) when compared to control (2.833±0.307 & 3.000±0.365) P value is < 0.001 & p<0.001. Dipping increased in RS+T (1.667±0.333 & 1.333±0.211) and p value is p<0.01 & p<0.05 group when compared to RS **Table 4**.

**Table 4.** Mean± SEM for Hole Board Test, Where DIP is dipping, SNI is sniffing, REA is rearing.

| GROUPS | CON | RS | RS+T | T |
| --- | --- | --- | --- | --- |
| PND42(DIP) | $2.833 \pm 0.307$ | $0.333 \pm 0.211$ | $1.667 \pm 0.333$ | $2.333 \pm 0.211$ |
| PND65(DIP) | $3.000 \pm 0.365$ | $0.333 \pm 0.211$ | $1.333 \pm 0.211$ | $2.333 \pm 0.211$ |
| PND42(SNI) | $1.000 \pm 0.258$ | $0.333 \pm 0.211$ | $0.667 \pm 0.333$ | $1.000 \pm 0.258$ |
| PND65(SNI) | $1.167 \pm 0.307$ | $0.167 \pm 0.211$ | $0.667 \pm 0.211$ | $1.167 \pm 0.307$ |
| **PND42(REA)** | 9.833±1.447 | 1.333±0.422 | 3.500±0.428 | 8.000±0.577 |
| **PND65(REA)** | 11.500±1.118 | 1.000±0.258 | 3.667±0.333 | 7.167±0.833 |

Sniffing was markedly varying among the groups. The decreased number of sniffing was observed in RS PND42 & PND65 (3.333±0.211 & 0.167±0.211) when compared to control (2.833±0.307 & 3.000±0.365) P value is >0.05 & P<0.05. Sniffing increased in RS+T (1.667±0.333 & 1.333±0.211) and p value is p>0.05 & p>0.05 group when compared to RS **Table 4**.

Rearing was markedly varying among the groups. The decreased number of sniffing was observed in RS PND42 & PND65 (1.333±0.422 & 1.000±0.258) when compared to control (2.833±0.307 & 3.000±0.365) P value is < 0.001 & P<0.001. Rearing increased in RS+T (9.833±1.447 & 11.500±1.118) and p value is p>0.05 & p<0.05 group when compared to RS **Table 4**.

Number of open arm entries was markedly varying among the groups. The decreased number of open arm entries was observed in RS PND42 & PND65 (2.677±0.333 & 3.667±0.422) when compared to control (5.333±0.667 & 8.000±0.577) P value is < 0.01 & P<0.001.open arm entries increased in RS+T (5.500±0.428 & 5.500±0.428) and p value is p<0.001 & p<0.05 group when compared to RS **Table 5**.

**Table 5.** Mean± SEM for Elevated plus maze where OAE is open arm entry, CAE is closed arm entry, HD is head dipping.

| **GROUPS** | **CON** | **RS** | **RS+T** | **T** |
| --- | --- | --- | --- | --- |
| **PND42(OAE)** | 5.333±0.667 | 2.677±0.333 | 5.500±0.428 | 5.167±0.601 |
| **PND65(OAE)** | 8.000±0.577 | 3.667±0.422 | 5.500±0.428 | 6.833±0.477 |
| **PND42(CAE)** | 3.333±0.333 | 6.500±0.428 | 4.167±0.601 | 3.667±0.333 |
| **PND65(CAE)** | 5.333±0.422 | 6.000±0.577 | 4.833±0.307 | 6.667±0.422 |
| **PND42(HD)** | 8.333±0.333 | 4.667±0.333 | 6.500±0.224 | 7.500±0.428 |
| **PND65(HD)** | 8.500±0.428 | 3.167±0.307 | 6.500±0.224 | 7.833±0.307 |

Number of closed arm entries was markedly varying among the groups. The increased number of open arm entries was observed in RS PND42 & PND65 (6.500±0.428 & 6.000±0.577) when compared to control (3.333±0.333 & 5.333±0.422) P value is < 0.001 & P>0.05.closed arm entries decreased in RS+T (4.167±0.601 & 4.833±0.307) and p value is p<0.01 & p<0.05 group when compared to RS **Table 5**.

Dipping was markedly varying among the groups. The decreased number of dipping was observed in RS PND42 & PND65 (4.667±0.333 & 3.167±0.307) when compared to control (8.333±0.333 & 8.500±0.428) P value is < 0.001 & P<0.001. Dipping increased in RS+T (6.500±0.224 & 6.500±0.224) and p value is p<0.001 & p<0.001 group when compared to RS **Table 5**.

Time spent with familiar animal was markedly varying among the groups. The increased time spent was observed in RS PND42 & PND65 (376.833±8.187 & 365.000±6.831) when compared to control (249.167±2.903 & 245.833±3.005) P value is < 0.001 & P<0.001. time taken decreased in RS+T (346.000±8.083 & 340.833±5.540) and p value is p<0.01 & p<0.05 group when compared to RS **Table 6**.

**Table 6.** Mean± Three chamber test. where FA is familiar animal, UFA is unfamiliar animal.

| GROUPS | CON | RS | RS+T | T |
| --- | --- | --- | --- | --- |
| PND42(FA) | 249.167±2.903 | 376.833±8.187 | 346.000±8.083 | 260.833±2.400 |
| PND65(FA) | 245.833±3.005 | 365.000±6.831 | 340.833±5.540 | 246.667±9.369 |
| PND42(UFA) | 320.500±8.086 | 207.667±14.186 | 243.000±22.283 | 286.667±13.556 |
| PND65(UFA) | 322.333±9.091 | 210.000±6.164 | 247.667±20.348 | 318.167±9.478 |

Time spent with unfamiliar animal was markedly varying among the groups. The decreased time spent was observed in RS PND42 & PND65 (207.667±14.186 & 210.000±6.164) when compared to control (320.500±8.086 & 322.333±9.091) P value is < 0.001 & P<0.001. time taken increased in RS+T (243.000±22.283 & 247.667±20.348) and p value is p>0.05 & p>0.05 group when compared to RS **Table 6**.

Number of central zone entries was markedly varying among the groups. The decreased number of entries was observed in RS PND42 & PND65 (0.667±0.333 & 0.333±0.211) when compared to control (8.000±0.856 & 9.000±0.966) P value is < 0.001 & P<0.001.number of entries increased in RS+T (1.500±0.428 & 0.667±0.211) and p value is p<0.05 & p<0.05 group when compared to RS **Table 7**.

**Table 7.**
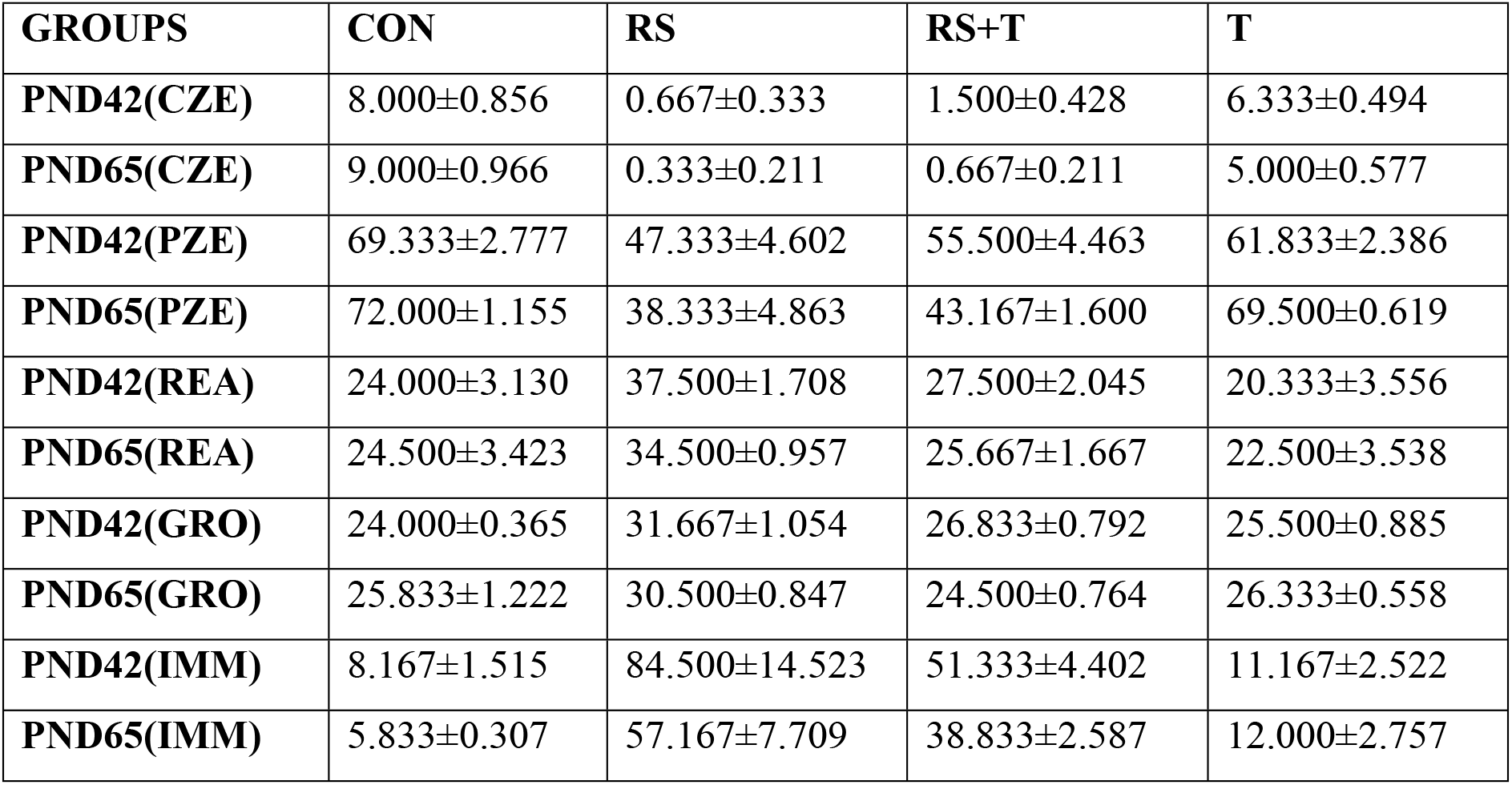
Mean± Open Field Task, where CZE is central zone entries is peripheral zone entries, REA is rearing, GRO is grooming and IMM is immobilization.

| GROUPS | CON | RS | RS+T | T |
| --- | --- | --- | --- | --- |
| PND42(CZE) | $8.000 \pm 0.856$ | $0.667 \pm 0.333$ | $1.500 \pm 0.428$ | $6.333 \pm 0.494$ |
| PND65(CZE) | $9.000 \pm 0.966$ | $0.333 \pm 0.211$ | $0.667 \pm 0.211$ | $5.000 \pm 0.577$ |
| PND42(PZE) | $69.333 \pm 2.777$ | $47.333 \pm 4.602$ | $55.500 \pm 4.463$ | $61.833 \pm 2.386$ |
| PND65(PZE) | $72.000 \pm 1.155$ | $38.333 \pm 4.863$ | $43.167 \pm 1.600$ | $69.500 \pm 0.619$ |
| PND42(REA) | $24.000 \pm 3.130$ | $37.500 \pm 1.708$ | $27.500 \pm 2.045$ | $20.333 \pm 3.556$ |
| PND65(REA) | $24.500 \pm 3.423$ | $34.500 \pm 0.957$ | $25.667 \pm 1.667$ | $22.500 \pm 3.538$ |
| PND42(GRO) | $24.000 \pm 0.365$ | $31.667 \pm 1.054$ | $26.833 \pm 0.792$ | $25.500 \pm 0.885$ |
| PND65(GRO) | $25.833 \pm 1.222$ | $30.500 \pm 0.847$ | $24.500 \pm 0.764$ | $26.333 \pm 0.558$ |
| PND42(IMM) | $8.167 \pm 1.515$ | $84.500 \pm 14.523$ | $51.333 \pm 4.402$ | $11.167 \pm 2.522$ |
| PND65(IMM) | $5.833 \pm 0.307$ | $57.167 \pm 7.709$ | $38.833 \pm 2.587$ | $12.000 \pm 2.757$ |

Number of peripheral zone entries was markedly varying among the groups. The decreased number of entries was observed in RS PND42 & PND65 (47.333±4.602 & 38.333±4.863) when compared to control (69.333±2.777 & 72.000±1.155) P value is < 0.001 & P<0.001.number of entries increased in RS+T (55.500±4.463 & 43.167±1.600) and p value is p>0.05 & p>0.05 group when compared to RS **Table 7**.

Rearing was markedly varying among the groups. The increased number of rearing was observed in RS PND42 & PND65 (37.500±1.708 & 34.500±0.957) when compared to control (24.000±3.130 & 24.500±3.423) P value is < 0.01 & P<0.05. Rearing decreased in RS+T (27.500±2.045 & 25.667±1.667) and p value is p<0.05 & p<0.05 group when compared to RS **Table 7**.

Grooming was markedly varying among the groups. The increased number of grooming was observed in RS PND42 & PND65 (31.667±1.054 & 30.500±0.847) when compared to control (24.000±3.130 & 25.833±1.222) P value is < 0.001 & P<0.001. grooming decreased in RS+T (26.833±0.792 & 24.500±0.764) and p value is p<0.001 & p<0.001 group when compared to RS **Table 7**.

Immobilization was markedly varying among the groups. The increased number of immobilizations was observed in RS PND42 & PND65 (84.500±14.523 & 57.167±7.709) when compared to control (8.167±1.515 & 5.833±0.307) P value is < 0.001 & P<0.001. immobilization decreased in RS+T (51.333±4.402 & 38.833±2.587) and p value is p<0.01 & p>0.05 group when compared to RS **Table 7**.

### Attention set shifting task

In **Simple discrimination** the number of trials is more was in PPS PND42 & PND65 (10.833±0.477 & 10.167±0.601) when compared to control (8.000±0.258 & 7.000±0.365) P value is < 0.001 & P<0.001. Trials are less in PPS+T (8.167±0.307 & 8.000±0.365) and p value is p<0.001& p<0.01 group when compared to PPS. P value P>0.05 is perceived in CON, PPS+TRT and TRT where as it is p<0.001 is seen in PPS+TRT in both TRT and VEC (7.500±0.428 & 6.500±0.563) and (7.333±0.211 & 6.500±0.428).

| GROUPS | CON | PPS | PPS+T | TRT ALONE | VEC |
| --- | --- | --- | --- | --- | --- |
| PND42 | $8.000 \pm 0.258$ | $10.833 \pm 0.477$ | $8.167 \pm 0.307$ | $7.500 \pm 0.428$ | $7.333 \pm 0.211$ |
| PND65 | $7.000 \pm 0.365$ | $10.167 \pm 0.601$ | $8.000 \pm 0.365$ | $6.500 \pm 0.563$ | $6.500 \pm 0.428$ |

In **Compound discrimination** number of trials increased in PPS PND42 & PND65 (10.833±0.477 & 9.333±0.333601) when compared to control (10.833±0.477 & 9.333±0.333)P value is < 0.001 & P<0.001. Trials are decreased in PPS+T (8.167±0.307 & 6.833±0.307) and p value is p<0.001 & p<0.01 group when compared to PPS. T PND42-PND65 (7.333±0.422 & 5.667±0.558) is significantly is appreciated where P < 0.001 only when compared to PP. Same way VEC PND42-PND65 (7.167±0.307&5.500±0.563) P>0.05 is seen in CON, PPS+T, TRT.

| GROUPS | CON | PPS | PPS+T | TRT ALONE | VEC |
| --- | --- | --- | --- | --- | --- |
| PND42 | $7.667 \pm 0.333$ | $10.833 \pm 0.477$ | $8.167 \pm 0.307$ | $7.333 \pm 0.422$ | $7.167 \pm 0.307$ |
| PND65 | $6.333 \pm 0.422$ | $9.333 \pm 0.333$ | $6.833 \pm 0.307$ | $5.667 \pm 0.558$ | $5.500 \pm 0.563$ |

Number of trials is increased in PPS PND42 & PND65 (12.167±0.477 & 10.833±0.477) when compared to control (7.667±0.333 & 6.667±0.2113) P value is < 0.001 & P<0.001. Trials are decreased in PPS+T (7.167±0.307 & 8.167±0.365) and p value is p<0.001 & p<0.01 group when compared to PPS in **Intradimenstional parameter** T PND42-PND65 (5,833±0.307 & 7.333±0.422) is significantly is appreciated where P < 0.001 only when compared to PP. Same way VEC PND42-PND65 (6.167±0.307 & 7.167±0.307) P>0.05 is seen in CON, PPS+T.

| GROUPS | CON | PPS | PPS+T | TRT ALONE | VEC |
| --- | --- | --- | --- | --- | --- |
| PND42 | 7.667±0.333 | 12.167±0.477 | 7.167±0.307 | 5.833±0.307 | 6.167±0.307 |
| PND65 | 6.667±0.211 | 10.833±0.477 | 8.167±0.365 | 7.333±0.422 | 7.167±0.307 |

The elevated number of trials was observed in **REV1** PPS PND42 & PND65 (15.000±0.816 & 12.833±0.477) when compared to control (10.667±0.422 & 9.333±0.494)P value is < 0.001 & P<0.001. Trials are decreased in PPS+T (13.333±0.422 & 10.500±0.428) and p value is p<0.001& p<0.01 group when compared to PPS .P value P>0.05 is perceived in CON, PPS+TRT and TRT where as it is p<0.001 is seen in PPS+TRT in both TRT and VEC (9.833±0.307&10.333±0.333) and (9.667±0.333 &8.500±0.428) respectively.

| GROUPS | CON | PPS | PPS+T | TRT ALONE | VEC |
| --- | --- | --- | --- | --- | --- |
| PND42 | 10.667±0.422 | 15.000±0.816 | 13.333±0.422 | 9.833±0.307 | 9.667±0.333 |
| PND65 | 9.333±0.494 | 12.833±0.477 | 10.500±0.428 | 10.333±0.333 | 8.500±0.428 |

**Extradimensional** number of trials was upshot in PPS PND42 & PND65 (39.667±1.667 & 32.000±1.000) when compared to control (24.500±1.384 & 20.000±1.390)P value is < 0.001 & P<0.001. Trials are downshifted in PPS+T (24.333±1.476 & 23.000±0.775) and p value is p<0.001 & p<0.01 group when compared to PPS. T PND42-PND65 (1.382±24.333 & 1.108±21.33325.500±0.885 & 26.333±0.558) is significantly is appreciated where P < 0.001 only when compared to PP. Same way VEC PND42-PND65 (24.333±1.085 & 21.333±1.022) P>0.05 is seen in CON, PPS+T, TRT.

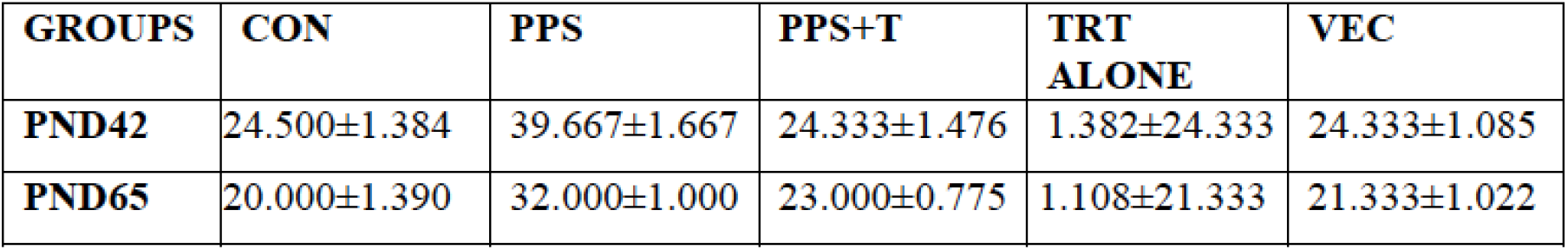

| GROUPS | CON | PPS | PPS+T | TRT ALONE | VEC |
| --- | --- | --- | --- | --- | --- |
| PND42 | 24.500±1.384 | 39.667±1.667 | 24.333±1.476 | 1.382±24.333 | 24.333±1.085 |
| PND65 | 20.000±1.390 | 32.000±1.000 | 23.000±0.775 | 1.108±21.333 | 21.333±1.022 |

In **REV2** upregulate the number of trials was observed in PPS PND42 & PND65 (31.667±0.615 & 23.000±0.816) when compared to control (23.000±0.775 & 19.333±0.422)P value is < 0.001 & P<0.001. Trials are decreased in PPS+T (24.333±1.476 & 19.333±0.667) and p value is p<0.001 & p<0.01 group when compared to PPS .P value P>0.05 is perceived in CON, PPS+TRT and TRT where as it is p<0.001 is seen in PPS+TRT in both TRT and VEC 20.500±0.957 & 18.333±0.615) and 20.500±0.619 & 19.000±0.447) respectively.

| GROUPS | CON | RS | RS+T | TRT<br>ALONE | VEC |
| --- | --- | --- | --- | --- | --- |
| PND42 | $23.000\pm0.775$ | $31.667\pm0.615$ | $24.333\pm1.476$ | $20.500\pm0.957$ | $20.500\pm0.619$ |
| PND65 | $19.333\pm0.422$ | $23.000\pm0.816$ | $19.333\pm0.667$ | $18.333\pm0.615$ | $19.000\pm0.447$ |

## DISCUSSION

Adolescence is categorized by heightened plasticity and behavioral flexibility[20].Spatial skills dependent upon neural structures that mature during adolescence, whereas exposure to chronic stress in adolescent disrupts the maturation of these structures, possibly negotiated in later part of life[21]. Experimental supports that mild to moderate level of stress early in life can alter HPA functions later in adulthood. For instance, both the rodents and primate studies, neonatal exposure to reoccurring bouts of novelty[22]. Repeated CRS leads to increase in number of spines and elongation of dendrites.

Evidence suggests that adolescent period hippocampal negative feedback to the HPA may be diminished. But in response to restraint stress in adolescent (PND 28) HPA showed longer latency to recover to baseline level of circulating ACTH and CORT [23]. Interestingly, human male and female adolescents(13-17 yrs.) also showed increased HPA activation in response to performance stressors exposure compared to male and female children (1-12 yrs.) and adults [24].

RAM is a standard and well validated test of spatial learning and memory. Working error are regarded as “short term” memory deficit and reference errors have been regarded as evidence of “long term” memory deficit [25]. Working memory error and reference memory error is significantly higher in adolescent RS compared to that of control of both PND 28 and PND 65. Where as in RS+ T(CA) it is significantly decreased compared to that of stress group. Time taken is also significantly higher in restraint group compared to that of control group **Fig 1**. Adolescents displayed higher activity levels in a novel environment, more rapidly approached a novel object in a familiar environment, and spent more time with a novel object relative to adults (Stanfield & Kirstein, et.al 2005). Object location and recognition is significantly reduced in RS group compared to control, where as in RS+T(CA) is significantly increased than the RS group **Fig 2**.

**Fig 1.**
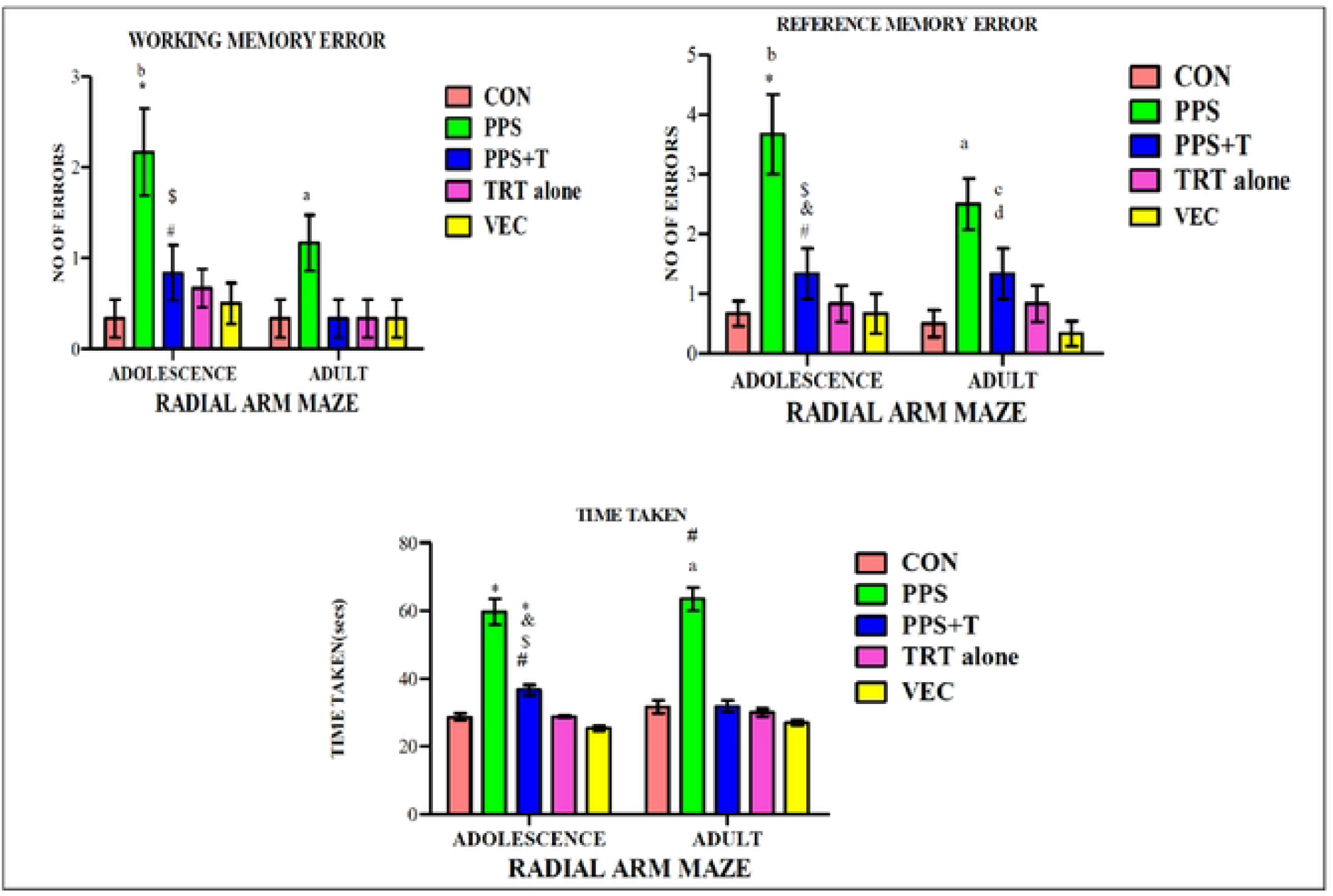
Radial arm maze.

**Fig 2.**
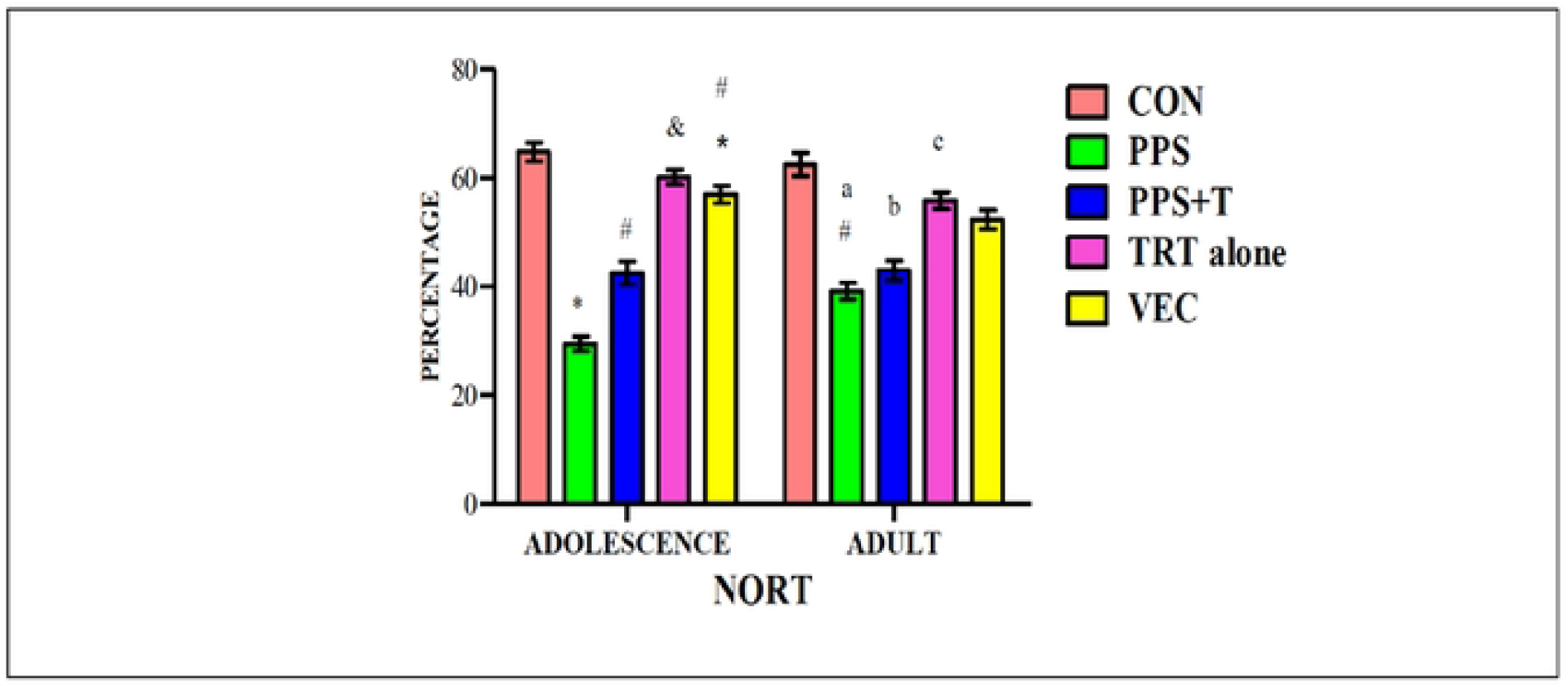
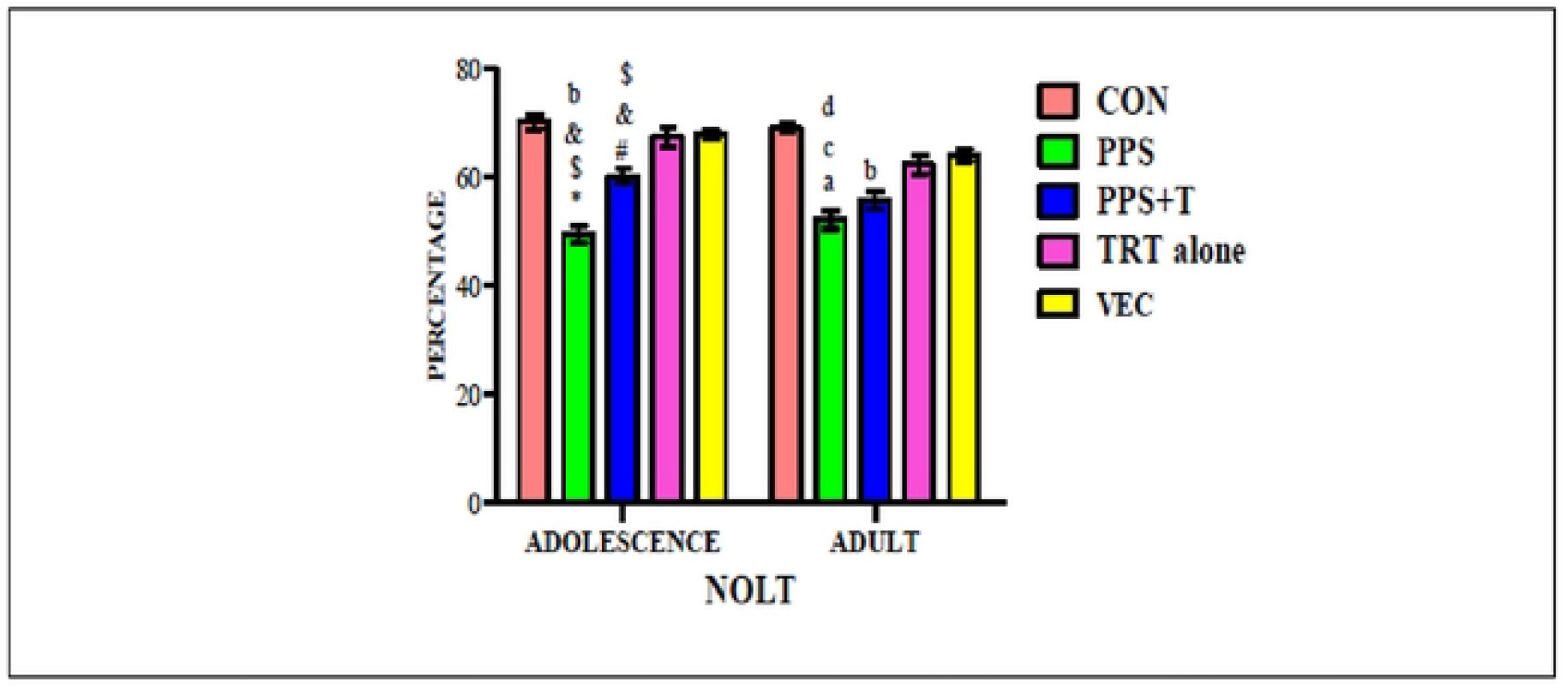
Novel object location task and Novel object recognition task.

The anxiolytic effect of stress was evidenced by the decrease in the exploration of open arms in EPM in stress rats when compared with the control group. Because the number of closed arm entries, which is an index of locomotor activity, was influenced, the decrease in the open arm exploration and therefore seems to be associated with the anxiolytic effects of stress in elevated plus maze. All measures are based on the principle of developing an avoidance approach conflict and indicate whether the animal follows its innate urge to explore new spaces, that is, the open arms or its fear of elevated, open spaces. RS showed decrease time spent in bright area and increase time spent in the dark area and number of fecal pellets. When treated with CA time spent in the bright area has been increased and decreased in time spent in dark area and number of fecal bolus **Fig 3**.

**Fig 3.**
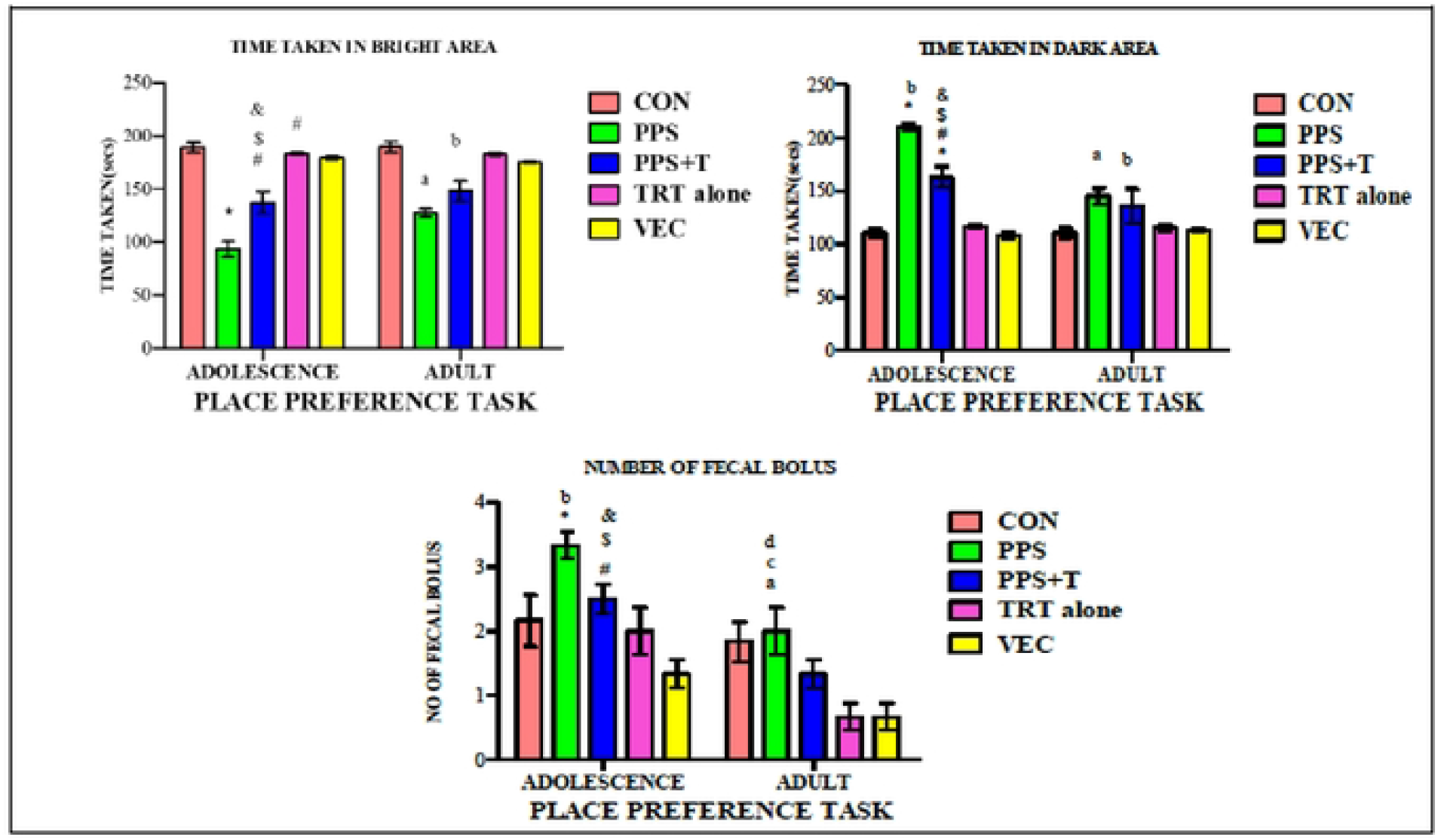
Place preference task.

Novel solution to a problem(task) have been reported to be related to novel exploratory behaviors, and a positive correlation [26]. Dipping, sniffing and rearing like behaviors are decreased in CRS compared to the control, whereas in RS+T significantly increased compared to the CRS in both PND42and PND65 **Fig 4**.

**Fig 4.**
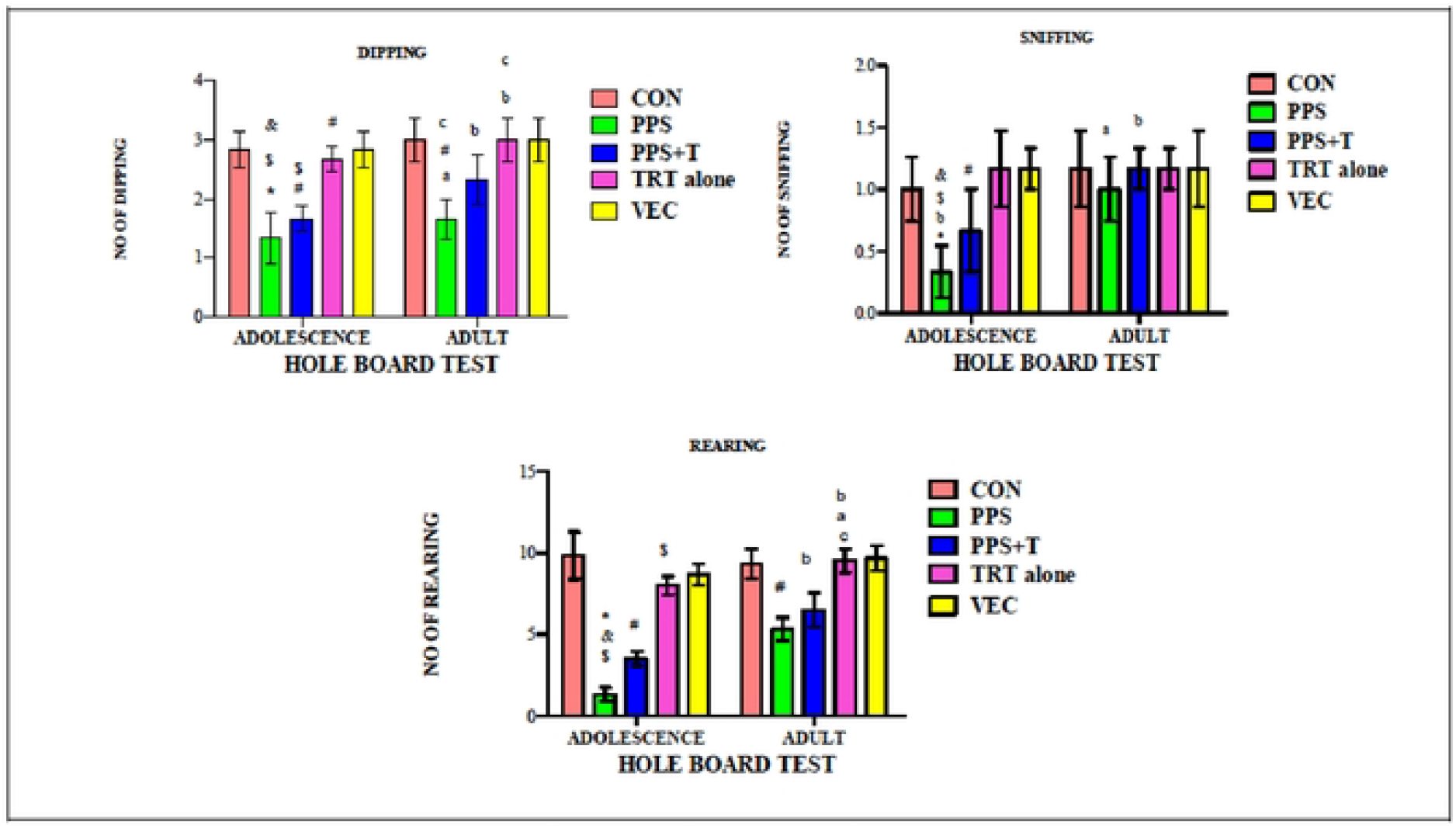
Hole Board Test.

In **Fig 5** Number of open arm entries and number of head dipping’s are decreased and number of closed arm entries and number of fecal bolus is increased compared to the control. whereas CRS+CA it is significantly increases the number of open arm entries and head dipping’s and decreases the closed arm entries and number of fecal bolus compared to the CRS.

**Fig 5.**
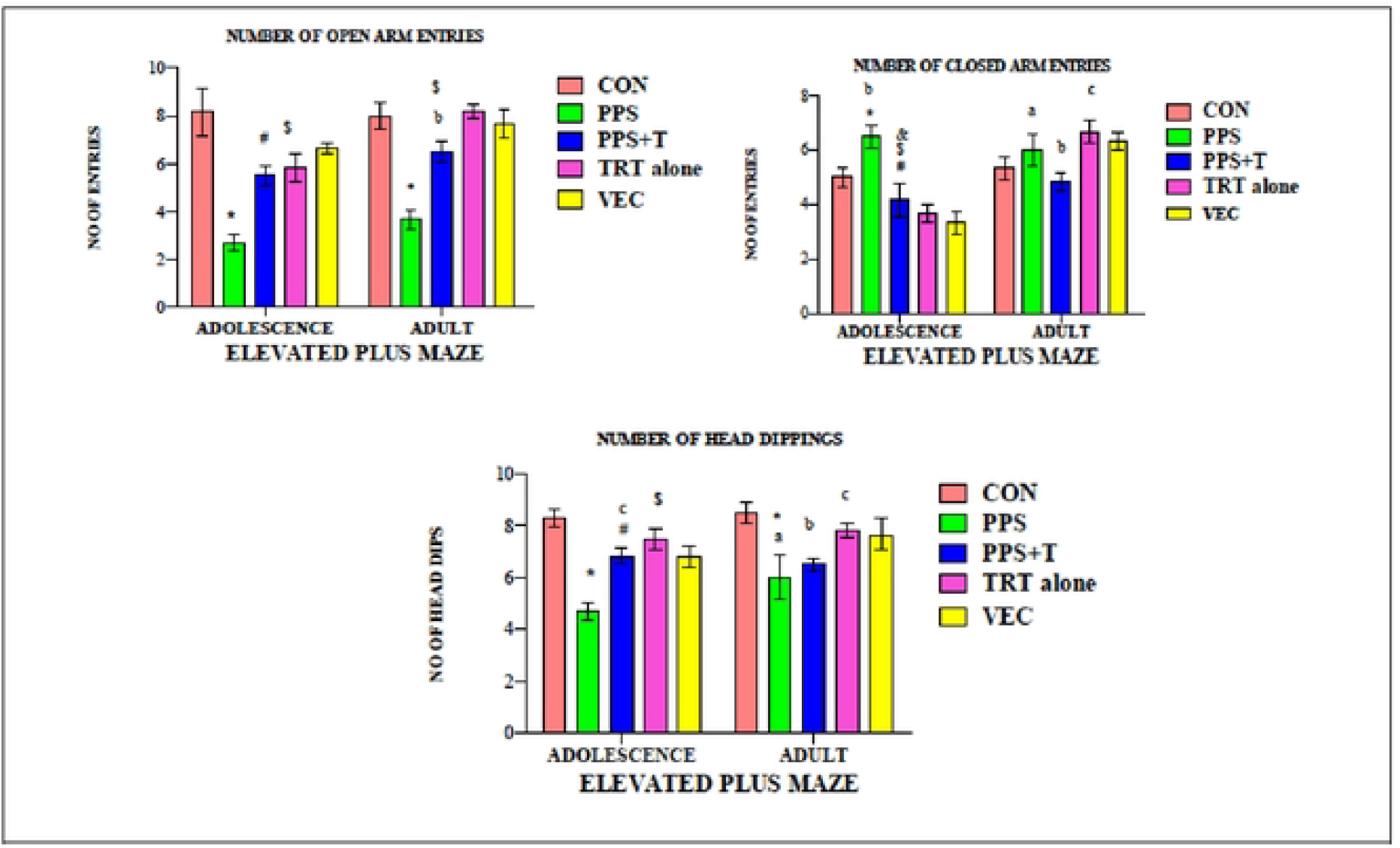
Elevated plus maze.

Stress being an important modulator of social behavior helps to deal with situation for survival and overcome threats by homeostasis. Chronic stress lowered social dominance as well as triggering depressive behavior in rodents[27] PND42 and PND65 CRS groups spent more time in familiar group compared to that of unfamiliar animal. CA+CRS significantly increases the social behavior to spent equal time with familiar and unfamiliar animal **Fig 6**.

**Fig 6.**
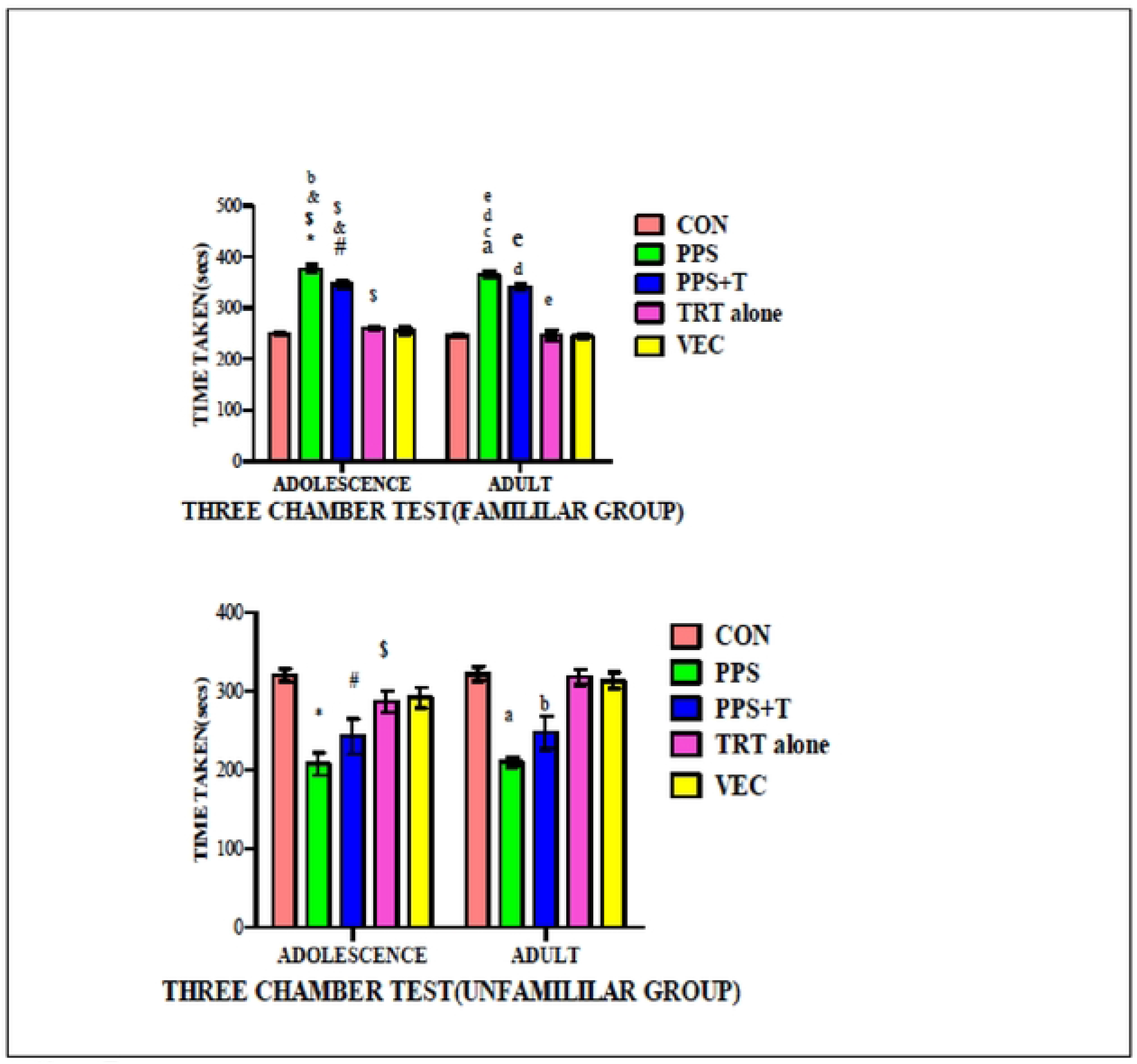
Three Chamber Test.

Decrease of exploratory behavior when the environment start to be familiar, [28] parameters can be also used (e.g., grooming, defecation, sitting) Contrary to habituation, sensitization is a non-associative learning in which the re-exposure to the initial stimulus increases the initial behavioral response. In CRS number of central zone entries and peripheral zone entries are significantly decreased and rearing, grooming and immobilization like behavior are significantly compared to the control group. But in RS+CA group number of central zone and peripheral zone entries is significantly increased and rearing, grooming and immobilization like behavior are significantly decreased compared to that of CRS group **Fig** 7.

**Fig 7.**
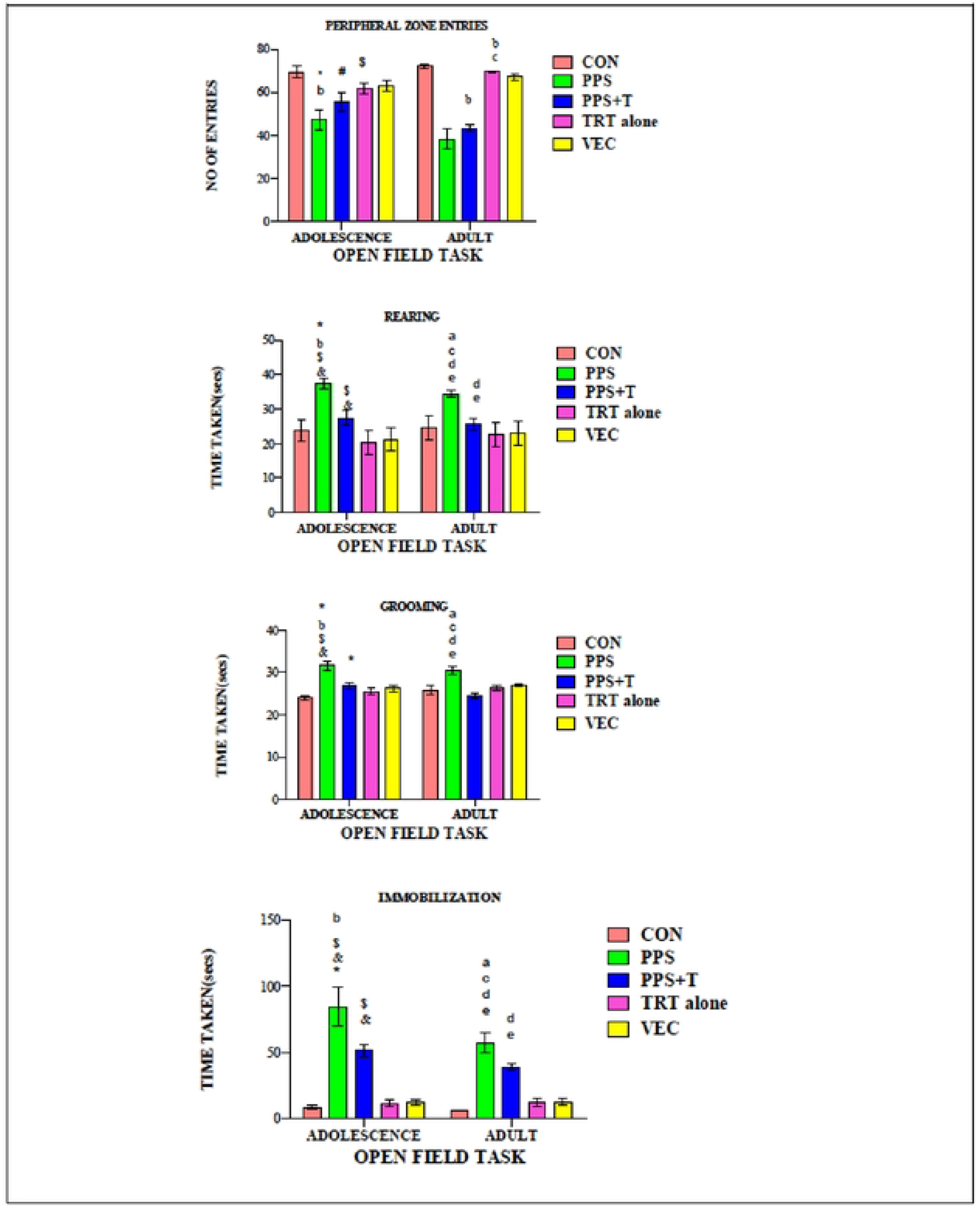
Open Fitld Task.

**Fig 8.**
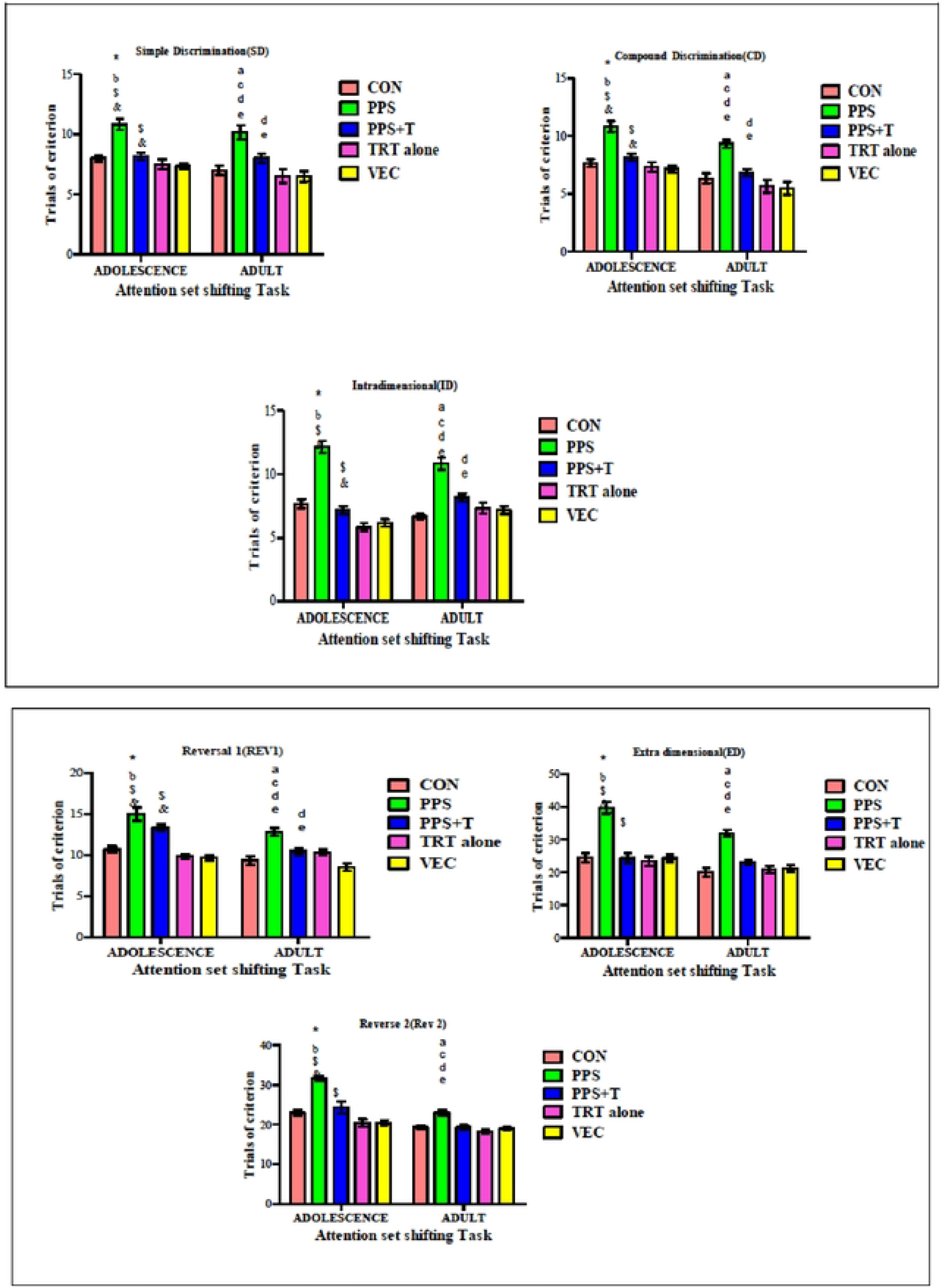
Attentional set shift task.

Distraction being a common behavior in adolescents compared to adults when exposed to irrelevant stimulus as part of complex stimulus. Adolescent rat younger than PND50 needs more trial than adults and result in lack of attentional flexibility. In all parameters such as SD, CD, ID, REV1, ED and REV2 adults rats performs better with less number of trials compared to the adolescence. Schizophrenic affected individuals has impaired reversal learning on set shifting

CA as an antioxidant and anti-inflammatory agent could improve learning and memory in stress group compared to the control group. Psychophysiological stress alters the pro-oxidant-antioxidant balance leads to mitochondrial dysfunction, disruption of energy pathways, neuronal damage, impaired neurogenesis and induction of signaling events in apoptotic cell death[29].

## Conclusion

The results showed that cinnamaldehyde treated rats exhibited restored neurobehavioral changes such as improved learning and memory, social behavior and decreased anxiety and depressive like behavior. CA may act as a promising drug like action for neurobehavioral deviations caused by stress.

